## Supplemental files for "Background optic flow modulates responses of multiple descending interneurons to object motion in locusts"

Table S1

| Animal | Total Number of Spikes | Number of Units | MANOVA (3D space) |
|---|---|---|---|
| L01 | 27214 | 16 | F(45,80793) = 38.426103, $p < 0.001$ |
| L02 | 25637 | 17 | F(48,76195) = 21.310296, $p < 0.001$ |
| L03 | 25383 | 18 | F(51,75511) = 47.208393, $p < 0.001$ |
| L04 | 30596 | 14 | F(39,90555) = 25.776592, $p < 0.001$ |
| L05 | 22838 | 15 | F(42,67699) = 22.374805, $p < 0.001$ |
| L06 | 23450 | 12 | F(33,69047) = 17.404464, $p < 0.001$ |
| L07 | 27229 | 20 | F(57,81123) = 68.932154, $p < 0.001$ |
| L08 | 27585 | 14 | F(39,81638) = 16.346922, $p < 0.001$ |
| L09 | 23615 | 14 | F(39,69882) = 25.803975, $p < 0.001$ |
| L10 | 24949 | 16 | F(45,74064) = 50.149337, $p < 0.001$ |
| L11 | 25347 | 17 | F(48,75333) = 45.143099, $p < 0.001$ |
| L12 | 24960 | 21 | F(60,74399) = 33.114825, $p < 0.001$ |
| L13 | 27171 | 22 | F(63,81034) = 26.620733, $p < 0.001$ |
| L14 | 21383 | 15 | F(42,63383) = 89.411021, $p < 0.001$ |
| L15 | 23769 | 19 | F(54,70760) = 48.159052, $p < 0.001$ |
| L16 | 31745 | 12 | F(33,93486) = 16.828258, $p < 0.001$ |
| L17 | 21400 | 11 | F(30,62776) = 10.113856, $p < 0.001$ |
| L18 | 26670 | 23 | F(66,79571) = 25.502046, $p < 0.001$ |
| L19 | 37880 | 19 | F(54,112806) = 95.960135, $p < 0.001$ |
| L20 | 38613 | 20 | F(57,105654) = 48.352062, $p < 0.001$ |
| L21 | 34081 | 17 | F(48,101310) = 78.549879, $p < 0.001$ |
| **Total** | 571515 | 352 | |
| **mean** | 27215 | 16.8 | |
| **median** | 25637 | 17 | |
| **s.d** | 4841.2 | 3.4 | |
| **min** | 21383 | 11 | |
| **max** | 38613 | 23 | |
| **1st quartile** | 23692 | 14 | |
| **3rd quartile** | 29090.5 | 19.5 | |

Table S2

| Animal | Total units | LW | LG | LF | F | None |
|---|---|---|---|---|---|---|
| L01 | 16 | 12 | 10 | 10 | 9 | 2 |
| L02 | 17 | 12 | 10 | 10 | 8 | 1 |
| L03 | 18 | 15 | 14 | 14 | 8 | 2 |
| L04 | 14 | 12 | 9 | 9 | 12 | 2 |
| L05 | 15 | 10 | 8 | 11 | 13 | 1 |
| L06 | 12 | 8 | 8 | 8 | 10 | 2 |
| L07 | 20 | 15 | 17 | 16 | 14 | 0 |
| L08 | 14 | 6 | 5 | 6 | 8 | 3 |
| L09 | 14 | 11 | 11 | 12 | 13 | 0 |
| L10 | 16 | 11 | 11 | 13 | 13 | 0 |
| L11 | 17 | 12 | 13 | 12 | 11 | 1 |
| L12 | 21 | 14 | 15 | 16 | 13 | 3 |
| L13 | 22 | 13 | 11 | 10 | 10 | 4 |
| L14 | 15 | 12 | 12 | 9 | 7 | 0 |
| L15 | 19 | 11 | 13 | 14 | 8 | 0 |
| L16 | 12 | 9 | 7 | 8 | 10 | 1 |
| L17 | 11 | 8 | 7 | 9 | 11 | 0 |
| L18 | 23 | 16 | 14 | 17 | 11 | 2 |
| L19 | 19 | 14 | 16 | 12 | 8 | 2 |
| L20 | 20 | 13 | 12 | 14 | 14 | 1 |
| L21 | 17 | 14 | 15 | 12 | 11 | 1 |
| **Total** | 352 | 248 | 238 | 242 | 222 | 28 |
| **mean** | 16.8 | 11.8 | 11.3 | 11.5 | 10.6 | 1.3 |
| **median** | 17 | 12 | 11 | 12 | 11 | 1 |
| **s.d.** | 3.4 | 2.6 | 3.2 | 2.9 | 2.2 | 1.2 |
| **min** | 11 | 6 | 5 | 6 | 7 | 0 |
| **max** | 23 | 16 | 17 | 17 | 14 | 4 |
| **1st quartile** | 14 | 10.5 | 8.5 | 9 | 8 | 0 |
| **3rd quartile** | 19.5 | 14 | 14 | 14 | 13 | 2 |

Table S3

| Response category | Number of units | | |
|---|---|---|---|
| | **LW** | **LG** | **LF** |
| Category 1: peak near TOC | 169 | 162 | 172 |
| Category 2: valley near TOC | 11 | 8 | 8 |
| Category 3: gradual increase | 21 | 27 | 5 |
| Category 4: tonic | 30 | 27 | 30 |
| Category 5: increase at the end | 17 | 14 | 27 |
| Total | 248 | 238 | 242 |

Table S4

| | LW | | LG | | LF | | F | |
|---|---|---|---|---|---|---|---|---|
| | **AIC** | **AICc** | **AIC** | **AICc** | **AIC** | **AICc** | **AIC** | **AICc** |
| **1 CT** | 24578.35 | 24590.03 | 23942.74 | 23953.95 | 24081.95 | 24093.11 | 117605.7 | 117608 |
| **2 CTs** | 22412.25 | 22459.59 | 22333.52 | 22378.96 | 21648.09 | 21693.33 | 114303.9 | 114312.8 |
| **3 CTs** | 21183.42 | 21291.85 | 21597.02 | 21701.06 | 20396.16 | 20499.76 | 118936.3 | 118956.5 |
| **4 CTs** | 21074.21 | 21270.73 | 20260.09 | 20448.62 | 19731.9 | 19919.63 | 114558.6 | 114594.5 |
| **5 CTs** | 19991.31 | 20304.66 | 19786.72 | 20087.25 | 18968.2 | 19267.44 | 111418.8 | 111474.9 |
| **6 CTs** | 19410.37 | 19871.14 | 19463.24 | 19905.06 | 18964.42 | 19404.35 | 111284.2 | 111364.9 |
| **7 CTs** | 19142.61 | 19783.47 | 19192.68 | 19807.06 | 18984.89 | 19596.63 | 111257.4 | 111367.3 |
| **8 CTs** | 19035.48 | 19891.37 | 19197.4 | 20017.74 | 19001.63 | 19818.42 | 111301 | 111444.5 |
